# Tensional homeostasis in multicellular clusters: effects of geometry and traction force dynamics

**DOI:** 10.1101/370437

**Authors:** J. Li, P. E. Barbone, M. L. Smith, D. Stamenović

## Abstract

The ability of cells to maintain a constant level of their cytoskeletal tension in response to external and internal disturbances is referred to as tensional homeostasis. It is essential for the normal physiological function of cells and tissues, and for protection against disease progression, including atherosclerosis and cancer. It has been shown recently that some cell types, such as endothelial cells, can maintain tensional homeostasis only when they form multicellular clusters, whereas other cell types, such as fibroblasts, do not require clustering for tensional homeostasis. For example, measurements of cell-extracellular matrix traction forces have shown that temporal fluctuations of the traction field in clusters of endothelial cells become progressively attenuated with increasing number of cells in the cluster, whereas in fibroblasts cell clustering does not influence traction field variability. Mechanisms that are responsible for these observations are largely unknown. In this study, a theoretical analysis and mathematical modeling have been applied to analyze experimental data obtained previously from traction microscopy measurements in order to investigate possible physical mechanisms that influence temporal variability of the traction field in multicellular forms. The focus of the analysis is on the contribution of dynamics and distribution of focal adhesion traction forces in conjunction with geometrical shape and size of multicellular clusters. Results of the analysis revealed that cluster size, magnitude and temporal fluctuations of focal adhesion traction forces have a major influence on traction field variability, whereas the influence of cluster shape appears to be minor.

## INTRODUCTION

Adherent cells exhibit the remarkable ability to adapt to applied mechanical stresses and strains. Because of this adaptation, cells can maintain their endogenous cytoskeletal mechanical tension at a steady and stable (homeostatic) level, which is essential for the normal physiological function of the tissues (1-8), and for protection against various diseases (2,3,9,10). While the idea of tensional homeostasis of cells was introduced two decades ago (11), there have been very few quantitative studies of this phenomenon. In 2004, Mizutani and colleagues (12) demonstrated that cellular stiffness returned to a set point level after stretching or relaxing single fibroblasts, which is an indirect indicator of tensional homeostasis in these cells. More recently, several additional quantitative studies of tensional homeostasis were reported. Webster and colleagues (13) have shown that in response to an applied step stretch, isolated fibroblasts do not return to the state of tension that they had prior to the stretch application. The authors referred to their observation as “tensional buffering” rather than tensional homeostasis. By contrast, we have observed that in isolated endothelial cells the traction field is not stable and exhibits large erratic temporal fluctuations (14,15), suggesting that individual endothelial cells cannot maintain tensional homeostasis. On the other hand, these fluctuations become attenuated in clusters of endothelial cells (15). These observations suggest that cell clustering may be required for stable (homeostatic) cytoskeletal tension in endothelial cells. In those studies, we introduced a first quantitative definition of tensional homeostasis as the ability of cells to maintain a consistent level of tension with low temporal fluctuations. It remains to be determined how, and indeed whether, tension is stabilized across multiple length scales, from focal adhesions to multicellular assemblies, and thus tensional homeostasis is itself a phenomenon that requires investigation across broad scales of both length and time.

One possible explanation for the observed attenuation of fluctuations of the traction field in clusters of endothelial cells is that it is a result of statistical averaging because of an increasing number of cells in a cluster. In this case, one would predict, based on the central limit theorem, that statistical averaging would lead to attenuation of fluctuations following an inverse square root dependence on the number of cells in the cluster, assuming traction fluctuations are independent of each other. Indeed, we have observed such dependence in non-confluent clusters. However, in confluent clusters, the rate of fluctuation attenuation is lower than the rate of the inverse square root dependence (15). Furthermore, in other cell types, such as fibroblasts, we have not observed a significant reduction of traction field fluctuations in multicellular clusters relative to single cells (16). Together, these observations suggest a possibility that factors other than statistical averaging may also affect traction field fluctuations in multicellular forms. In this study, we investigated how dynamics and distribution of focal adhesion traction forces within a cluster, as well as cluster shape and size affected traction field variability. Our rationale was as follows.

We and others have observed that traction force distribution within a cluster is not uniform. Forces of large magnitude are applied at focal adhesions (FAs) located near the cluster edges, whereas forces of low magnitudes are applied at FAs located in the cluster interior. For example, Maruthamuthu et al. (17) used traction force microscopy to measure cell-extracellular matrix (ECM) traction forces in two- and three-cell cluster of epithelial cells. They showed that large traction forces were distributed near the cluster outer boundary and that low traction forces were distributed in the cluster interior. Mertz et al. (18) measured cell-ECM traction forces in colonies of keratinocytes before, during, and after formation of cadherin-mediated intercellular adhesions. As cadherin-dependent junctions form, there was dramatic rearrangement of traction forces from a disorganized distribution to an organized concentration of force at the colony periphery. We used traction microscopy to measure cell-ECM traction forces in multicellular clusters of fibroblasts (19) and in multicellular clusters of endothelial cells (15). We observed that in both cell types large traction forces were distributed along the cluster edges and small traction forces were distributed in the cluster interior. We have also observed that temporal fluctuations of traction forces relative to their mean values decrease with increasing magnitude of those forces, regardless of the cell type or the cluster size (15,16). Taken together, all these observations suggest that variability of the traction field may depend on the shape and the size of the cluster in conjunction with variability of FA traction forces. First, with increasing cluster size one would expect that the number of interior forces, which are of smaller magnitude and more variable, would increase relative to the number of edge forces, which are of greater magnitude and less variable. Thus, with increasing cluster size, one might expect an increasing variability of the traction field, which is opposite from the effect of statistical averaging. Second, for clusters having the same projected area (or the same number of FAs), one would expect that non-convex-shaped clusters would have a larger number of edge forces relative to the interior forces than convex-shaped clusters because of the larger circumference-to-area ratio in the former than in the latter.

In this study we analyzed how magnitude, variability, and the numbers of edge and interior traction forces within a cluster affected variability of the overall traction field of the cluster. We combined this analysis with mathematical models of multicellular clusters of different sizes and shapes where we simulated traction force fluctuations using a Monte Carlo approach. Finally, we combined these models with experimental data for FA traction force fluctuations that we obtained previously from clusters of living endothelial cells and fibroblasts. Results of our study revealed that traction field variability was highly influenced by the magnitude, distribution, and dynamics of traction forces, and by cell cluster size. The influence of cluster shape, on the other hand, appeared to be minor.

## METHODS AND RESULTS

### Analysis of the Coefficient of Variation

We assumed that at a given time (*t*) there were two types of traction forces of different magnitudes acting on the cluster: larger edge forces, and smaller interior forces. We used a scalar metric of the traction field that we used before (15), namely the net traction force (*T*), which we defined as the sum of the magnitudes of all traction forces in the cluster at a given *t*, i.e.,

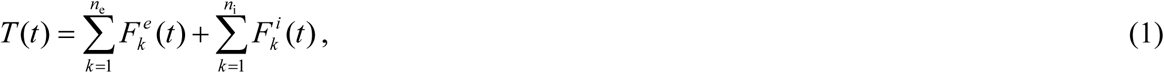

where *F^e^*(*t*) and *F^i^*(*t*) are magnitudes and, *n_e_* and *n_i_* are numbers of the edge and the interior forces at time *t*, respectively, such that *n* = *n_e_* + *n_i_*, is the total number of traction forces in the cluster.

Since in our previous studies we measured traction forces at discrete time intervals, the averages of *F^e^*(*t*) and *F^i^*(*t*) over time period τ, are 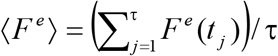 and 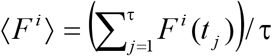, respectively. We assumed that *F^e^*(*t*) and *F^i^*(*t*) fluctuated around 〈*F^e^*〉 and 〈*F^i^*〉, which were the same for all edge and all internal forces, respectively. We also assumed that these fluctuations were not correlated, and that their corresponding variances were σ^2^ (*F^e^*) and σ^2^ (*F^i^*). Thus it follows from Eq. 1 that the time average of *T*(*t*) is

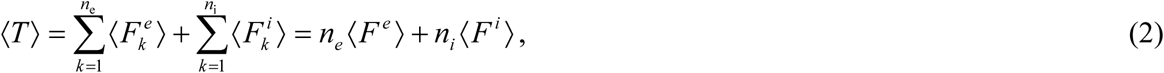

and that the variance [σ^2^ (*T*)] of *T*(*t*) is

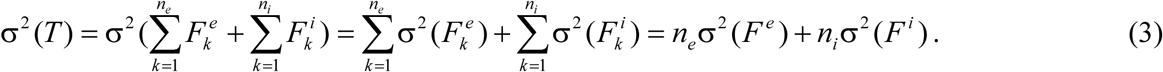

Using Eqs. 2 and 3, we obtained the coefficient of variation (*CV_T_*) of *T*(*t*) as follows

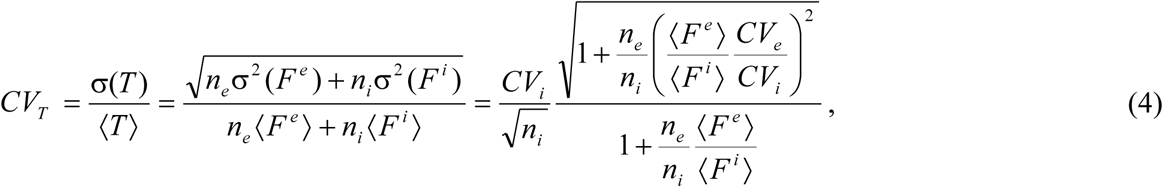

where *CV_e_* = σ(*F^e^*) /〈*F^e^*〉 and *CV_i_* = σ(*F^i^*) /〈*F^i^*〉 are the coefficients of variation of *F^e^*(*t*) and *F^i^*(*t*), respectively. Equation Eq. 4 shows that variability of the traction field depends on variability, magnitudes, and relative numbers of edge and interior traction forces.

Our previous traction measurements have shown that fluctuations of FA traction forces do not depend on the number of cells in the cluster and on the cell type (15,16). For single cells, as well as for multicellular cluster of various sizes and shapes, temporal fluctuations of FA traction force (i.e., their coefficient of variation *CV_F_*) decrease with increasing mean force (〈*F*〉). For single bovine aortic endothelial cells (BAECs) and for confluent clusters of BAECs of different sizes, the *CV_F_* vs. 〈*F*〉 relationship exhibits a weak, but significant negative correlation (Spearman correlation coefficient ρ = −0.244, *p* = 2×10^−7^) (Fig. 1). A similar, somewhat stronger correlation (ρ = −0.441, *p* = 2×10^−7^) has been obtained for clusters of mouse embryonic fibroblasts (MEFs) (Fig. S1 of the Supplemental Information).

**Figure 1.**
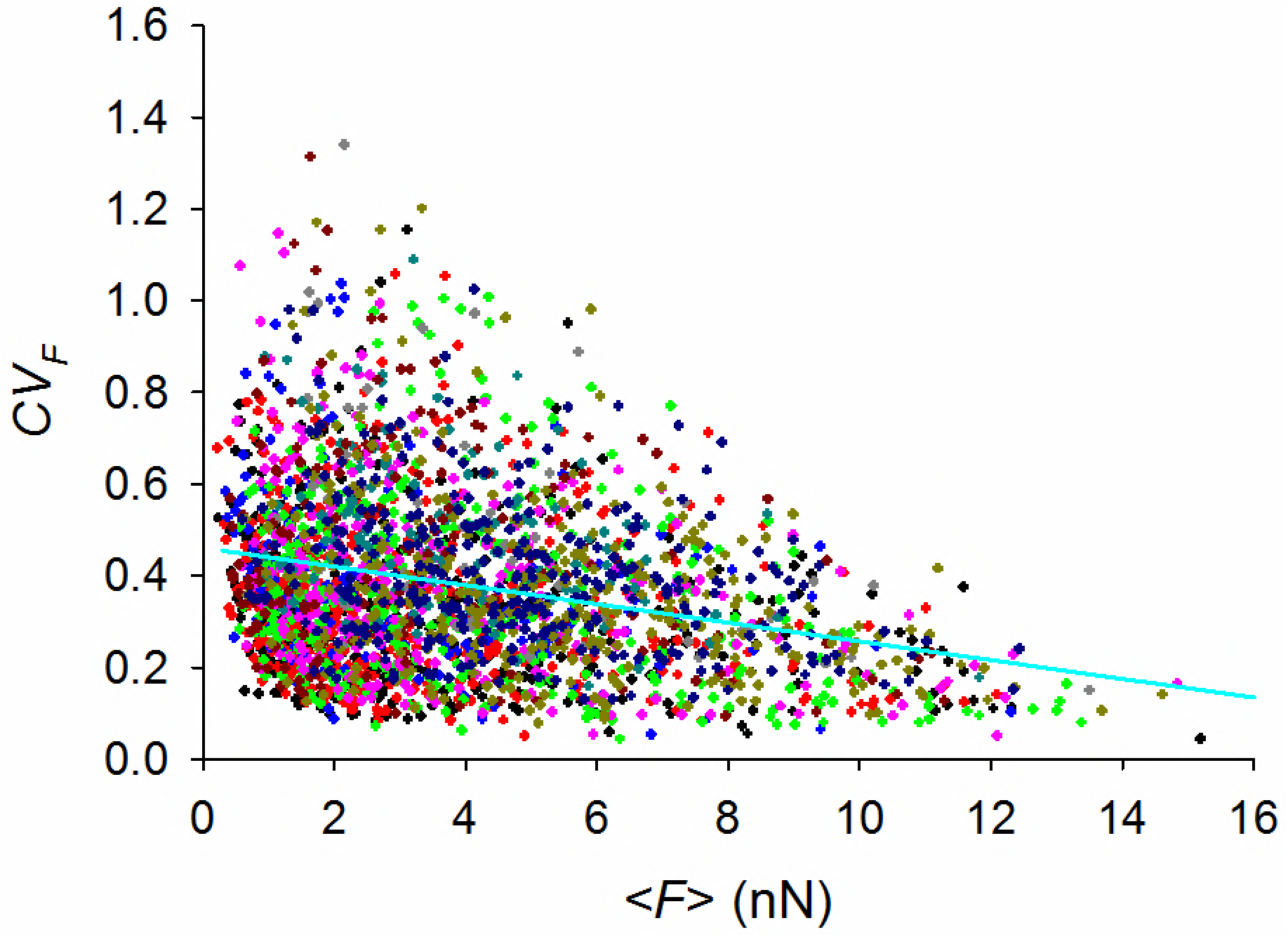
Coefficient of variation of FA forces (*CV_F_*) decreases with increasing of the mean force (〈*F*〉). Data are obtained from traction microscopy measurements in single BAECs and in confluent clusters of BAECs (15). Each color represents different cluster size ranging from single cells to 30-cell clusters. The cyan line is the linear regression best fit, *CV_F_* = 0.46 – 0.021 〈*F*〉.

Based on the data in Fig. 1, we assumed that *F^e^* and *F^i^* did not depend on *n_e_* and *n_i_* and that *CV_e_* and *CV_i_* had the same correlation with 〈*F^e^*〉 and 〈*F^i^*〉, respectively, as the data shown in Fig. 1. Taking these assumptions into account, it follows from Eq. 4 that *CVT* is a function of four independent variables *n_e_*, *n_i_*, 〈*F^e^*〉 and 〈*F^i^*〉. It is clear that for the case where 〈*F^e^*〉 = 〈*F^i^*〉, and therefore *CV_e_* = *CV_i_*, Eq. 4 becomes 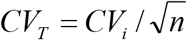, i.e., fluctuations of the traction field scale with fluctuations of individual traction forces according to an inverse square root relationship. For the case where 〈*F^e^*〉 > 〈*F^i^*〉, we graphed *CV_T_* as a function of *n_e_/n_i_* and 〈*F^e^*〉*/*〈*F^i^*〉, for different values of *n*, according to Eq. 4, using an empirical relationship *CV_F_* = 0.46 – 0.021〈*F*〉, obtained by fitting the data for BAECs from Fig. 1, and setting arbitrarily 〈*F^i^*〉 = 1 nN. The values of *n* corresponded to the average number of FAs observed previously (15) in single BAECs (*n*=18) and in clusters of BAECs ranging from 2-cell to 30-cell clusters (*n* = 27, 35, 46, 57, 73, 88, 109, 133, 164, 247, 402). We found that *CV_T_* became attenuated with increasing 〈*F^e^*〉*/*〈*F^i^*〉 and with decreasing *n_e_/n_i_* (Fig. 2A), and with increasing *n* (Fig. 2B).

To further investigate how cluster size, shape and dynamics of traction forces affect variability of the traction field we developed a more detailed mathematical model.

**Figure 2.**
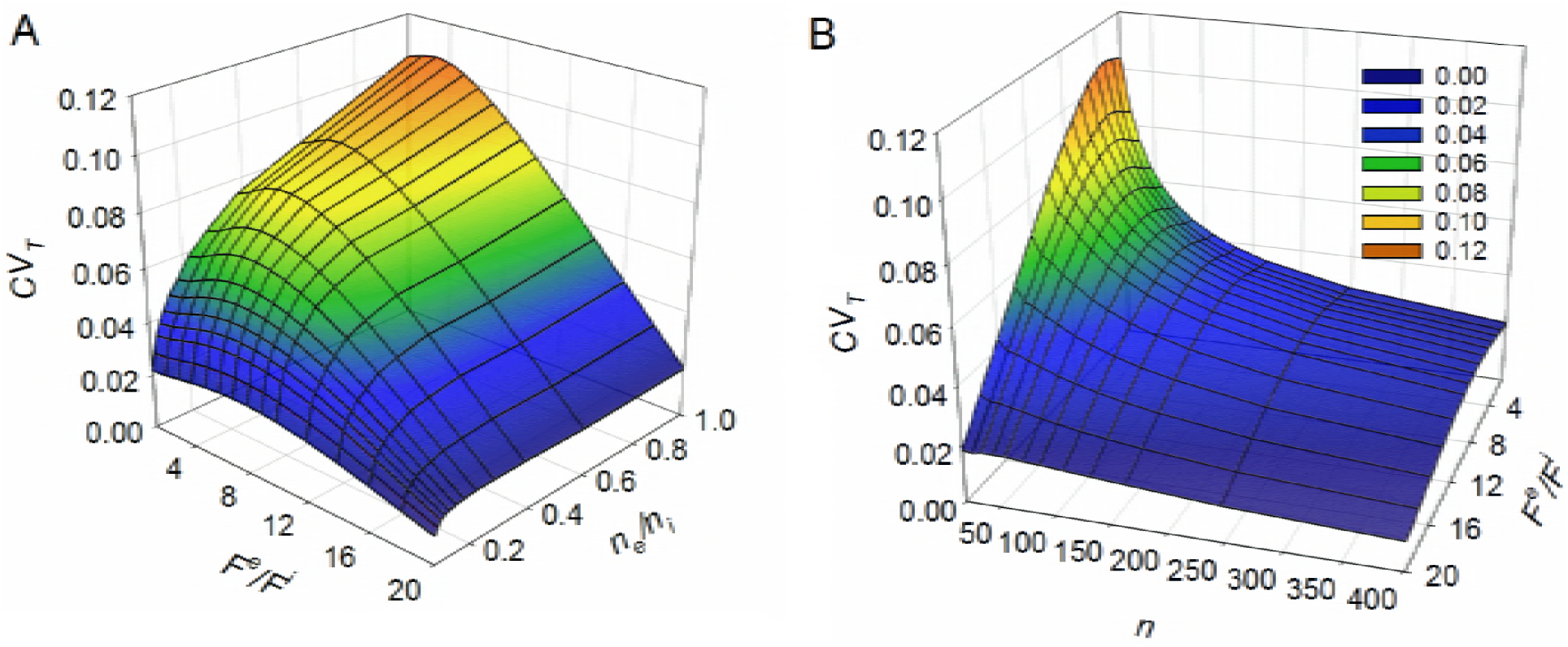
Variability of the traction field (*CV_T_*) as a function of *n_e_/n_i_* and *F^e^/F^i^* (A) and as a function of *n* and *F^e^/F^i^* (B). The graphs are obtained from Eq. 4 and the linear regression from Fig. 1.

### Mathematical Model

For the purpose of modeling, we considered tractions forces as externally applied forces. We modeled confluent cell clusters as two-dimensional arrays of massless unit square blocks (“cells”). Two types of clusters were considered: square-shaped (convex) clusters (Fig. 3A) and arbitrary shaped (non-convex) clusters (Fig. 3B) composed of *N* = 1, 4, 9, 16, 25, 36, 49, … 121 square cells. We applied 8 forces to each cell: four at the corners, and four smaller forces at the interior of each cell. Since measurements in living clusters show that forces near the edges are greater than the interior forces, we applied larger forces at cell corners that lie along the edge of the cell-cluster, and smaller forces to cell corners that are on the interior of the cell cluster. These forces are shown in a few example clusters in Fig. 4. All traction force vectors were directed along the diagonals of the cell, pointing away from the center of the cell.

**Figure 3.**
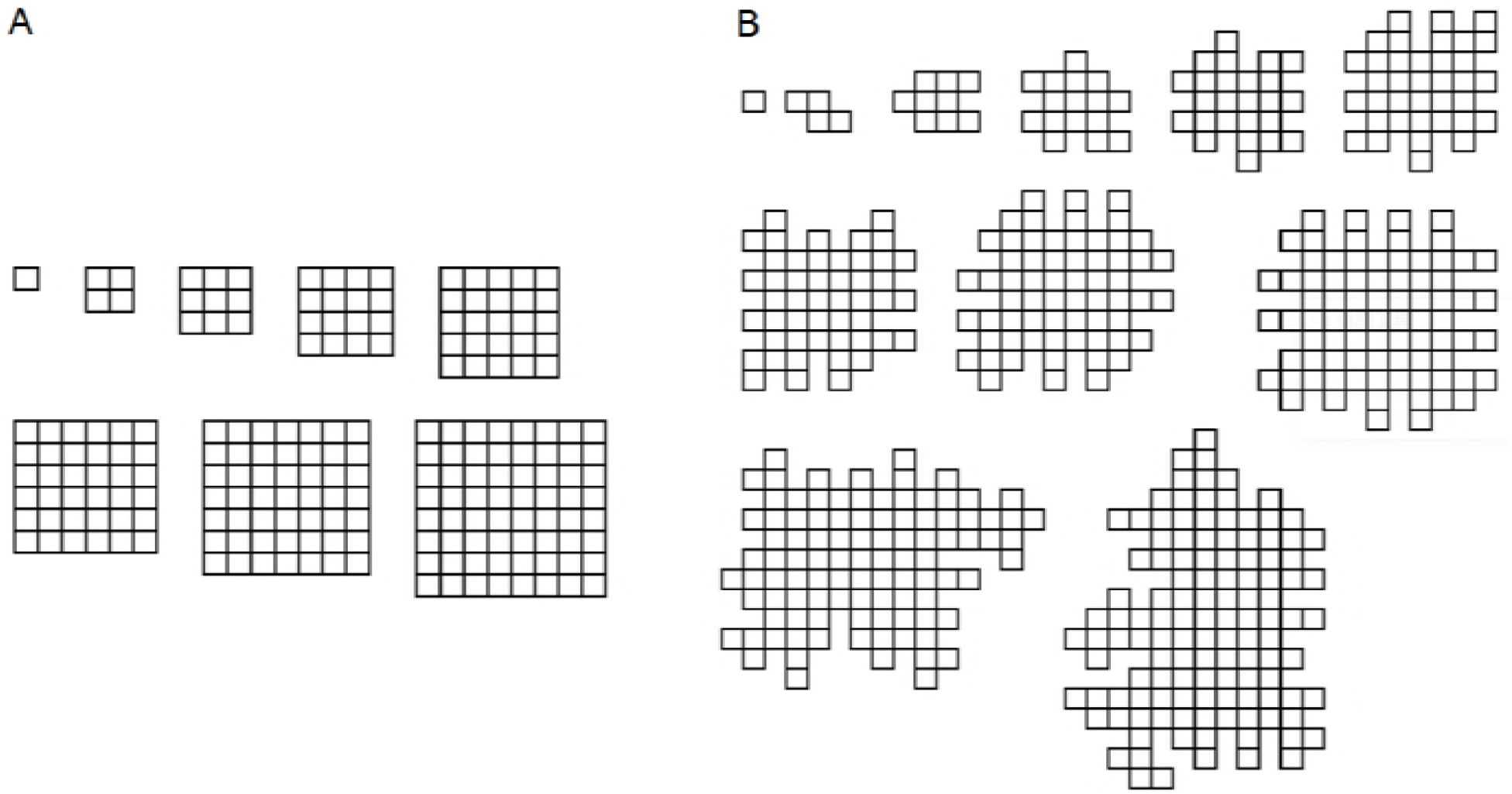
Models of square (convex) clusters (A) and of non-convex clusters (B). The number of cells in the clusters range from 1 to 121 cells.

For a given number of cells (*N*) in the cluster, the total number of edge forces in square clusters is 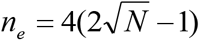 and the total number of interior forces is 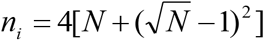. Fornon-convex clusters, corresponding values of *n_e_* and *n_i_* for a given *N* are given in Table 1. Note that for both square and non-convex clusters, the total number of forces in the cluster is *n* = *n_e_ + n_i_* = 8*N*. By combining the values of *ne* and *ni* from Table 1 with Eq. 4 and using the same *CV_F_* vs. 〈*F*〉 regression for BAECs (i.e., *CV_F_* = 0.46 – 0.021〈*F*〉), we obtained *CV_T_* vs. *N* relationship for square and non-convex clusters for 〈*F^e^*〉*/*〈*F^i^*〉 = 8 and 〈*F^e^*〉*/*〈*F^i^*〉 = 20 (Fig. 5). The graphs show that the higher the ratio 〈*F^e^*〉*/*〈*F^i^*〉 is, the more attenuated *CV_T_* is, and the slower the rate of attenuation of *CV_T_* with increasing *N* is. This rate of attenuation is higher in the non-convex than in the square clusters, but slower than the inverse square root dependence.

**Figure 5.**
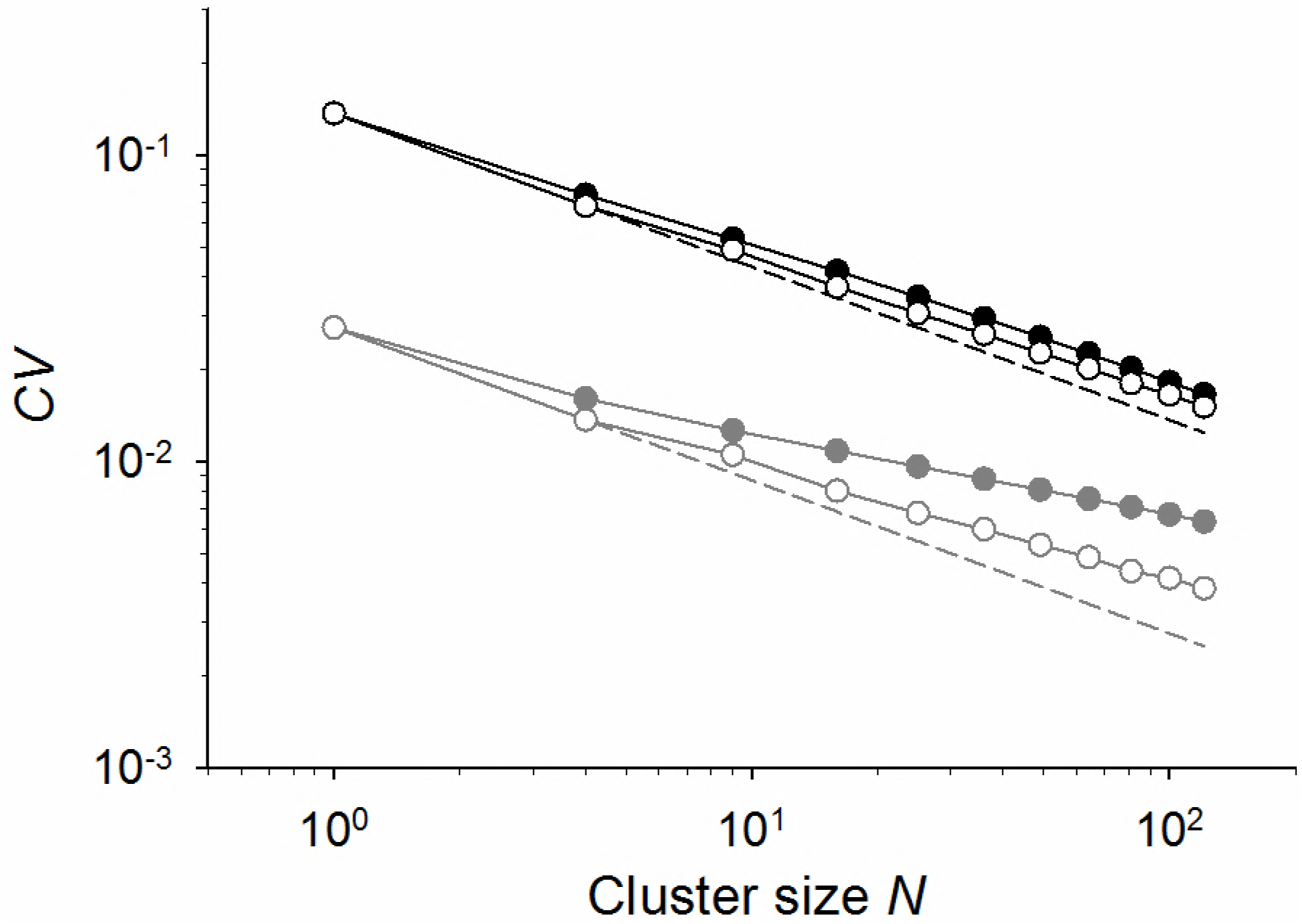
Coefficient of variation (*CV_T_*) vs. cluster size (*N*) for *F^e^/F^i^* = 8 (black) and *F^e^/F^i^* = 20 (gray) obtained from Eq. 4 and the linear regression fitted to the *CV_F_* vs 〈*F*〉 experimental data for BAEC cells and clusters (Fig. 1). The solid circles correspond to the square clusters and the open circles correspond to non-convex clusters. The dashed lines correspond to the inverse square root dependence, 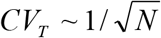.

**Table 1.**
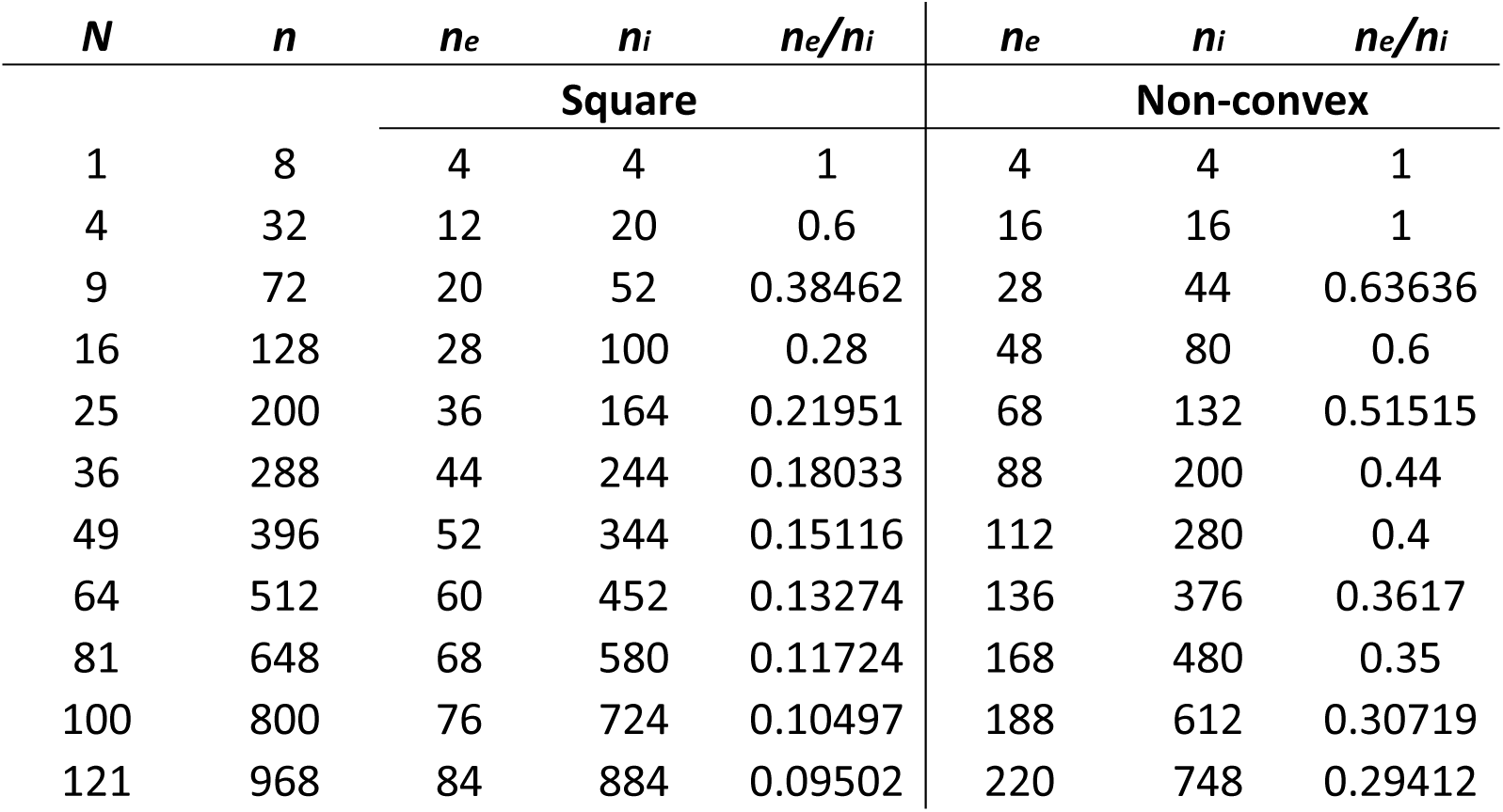
Values for the total number of cells (*N*), the total number of traction forces (*n*), the total number of edge forces (*n_e_*) and of interior forces (*n_i_*) for *n_e_*/*n_i_* ratio for square (convex) clusters and for non-convex clusters.

We repeated these calculations using a linear regression *CV_F_* =0.38 – 0.026 〈*F*〉 obtained from fitting *CVF* vs. 〈*F*〉 for MEFs. We obtained that for square clusters *CV_T_* had lower values and a lower rate of attenuation in MEFs than in BAECs (Fig. S2 of Supplemental Information).

#### Monte Carlo Simulations

We applied forces to the mathematical model to investigate how force dynamics affects variability of the traction field of the model. We used a Monte Carlo approach to simulate traction force fluctuations. Forces were fluctuating at 5 min time intervals over 2 h, which was consistent with force fluctuations observed experimentally (15). A description of the Monte Carlo algorithm is provided in Supplemental Information (Section S1).

Representative Monte Carlo force fluctuations are shown in Fig. 6A. For comparison, time lapses of measured physical forces of similar magnitudes from our previous study (15) are shown in Fig. 6B.

**Figure 6.**
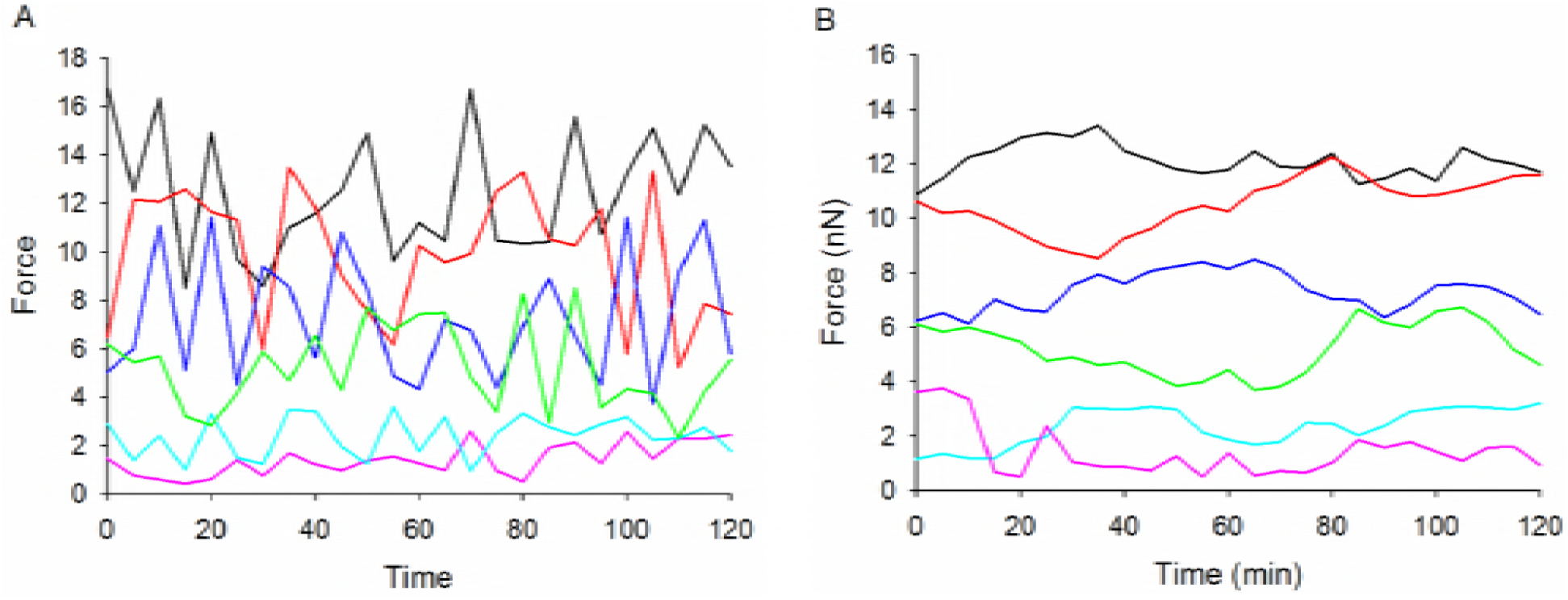
Representative examples of Monte Carlo force fluctuations (A) and experimental FA force fluctuations (B). Each color corresponds to the same mean force.

Monte Carlo forces also exhibited a negative correlation between *CV_F_* and 〈*F*〉 (ρ = −0.641, *p* = 2×10^−7^) (Fig. 7). Fitting this relationship with a linear regression yielded *CV_F_* = 0.47 – 0.022 〈*F*〉, which was very similar to the linear regression that was obtained for BAECs. This was expected since the Monte Carlo forces were generated based on the *CV_F_* vs. 〈*F*〉 relationship of BAECs FA forces (Fig. 1), and thus provides a check on the simulation.

**Figure 7.**
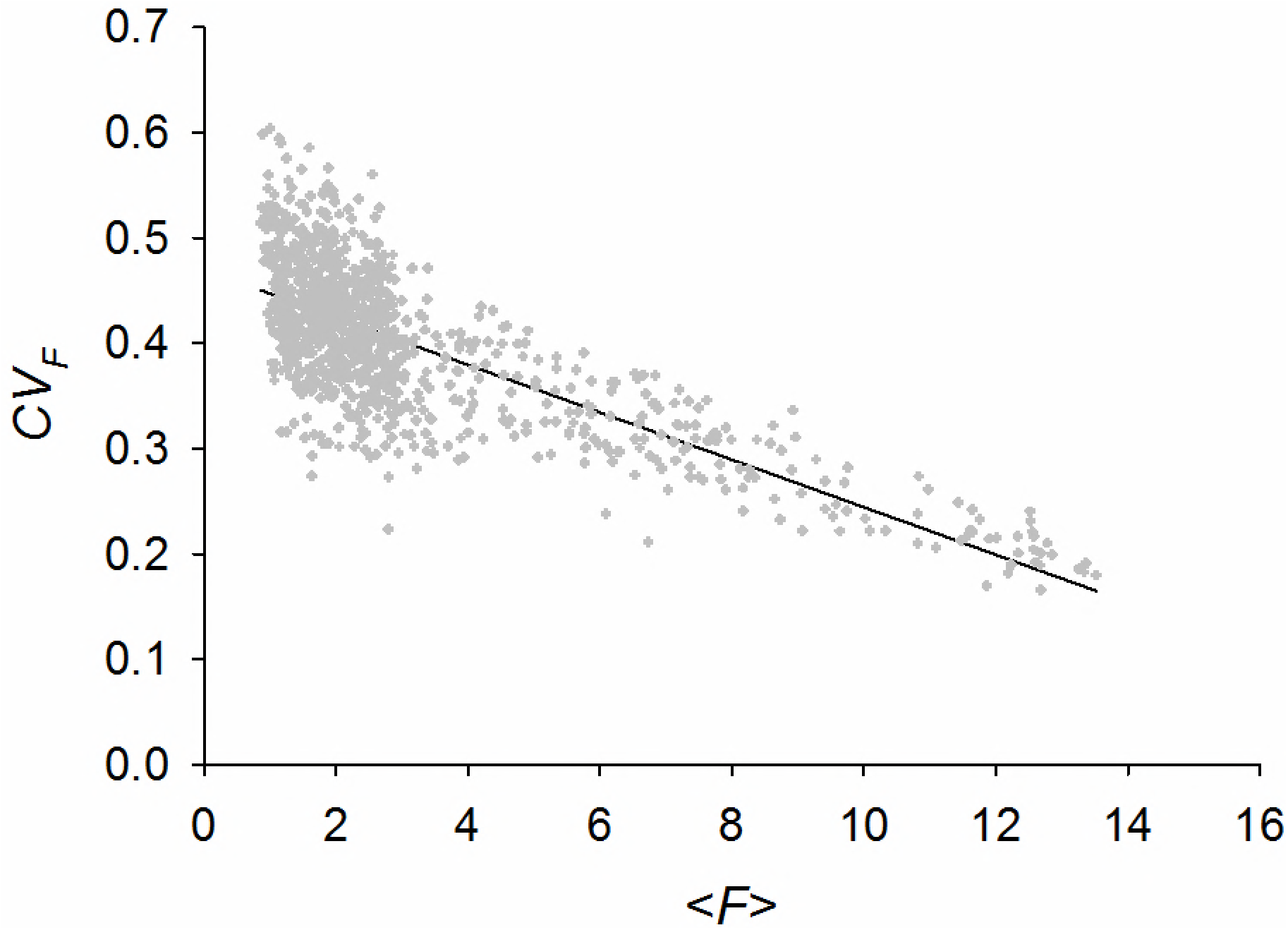
Coefficient of variation of Monte Carlo forces (*CV_F_*) decreases with increasing of the mean force (〈*F*〉). Data are obtained from Monte Carlo simulations of BAEC forces applied to the 121-cell cluster model. The black line is the linear regression best fit, *CV_F_* = 0.47 – 0.022〈*F*〉.

Initially, all forces were acting along the diagonals of the square unit, as shown in Fig. 4. In order to maintain equilibrium of a cluster at each time interval, force vectors were adjusted by a least squares procedure to the closest equilibrium configuration, as we did previously (20). After equilibration, we calculated at every 5-min interval the net traction force of a cluster as the sum of the norms of the Monte Carlo force vectors, i.e.,

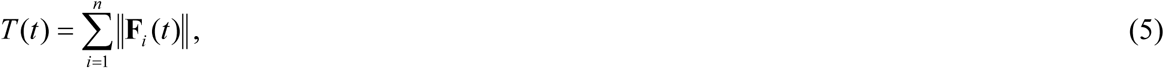

where *n* is the number of traction forces applied to a cluster. From the time lapses of *T*(*t*), we computed its time-average, the corresponding standard deviation and *CV_T_* as the ratio of the two
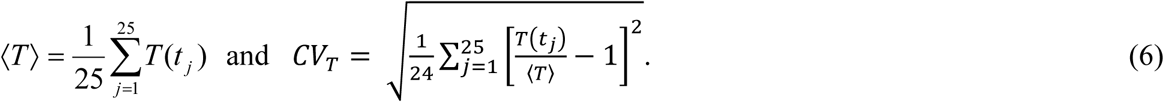

**Figure 4.**
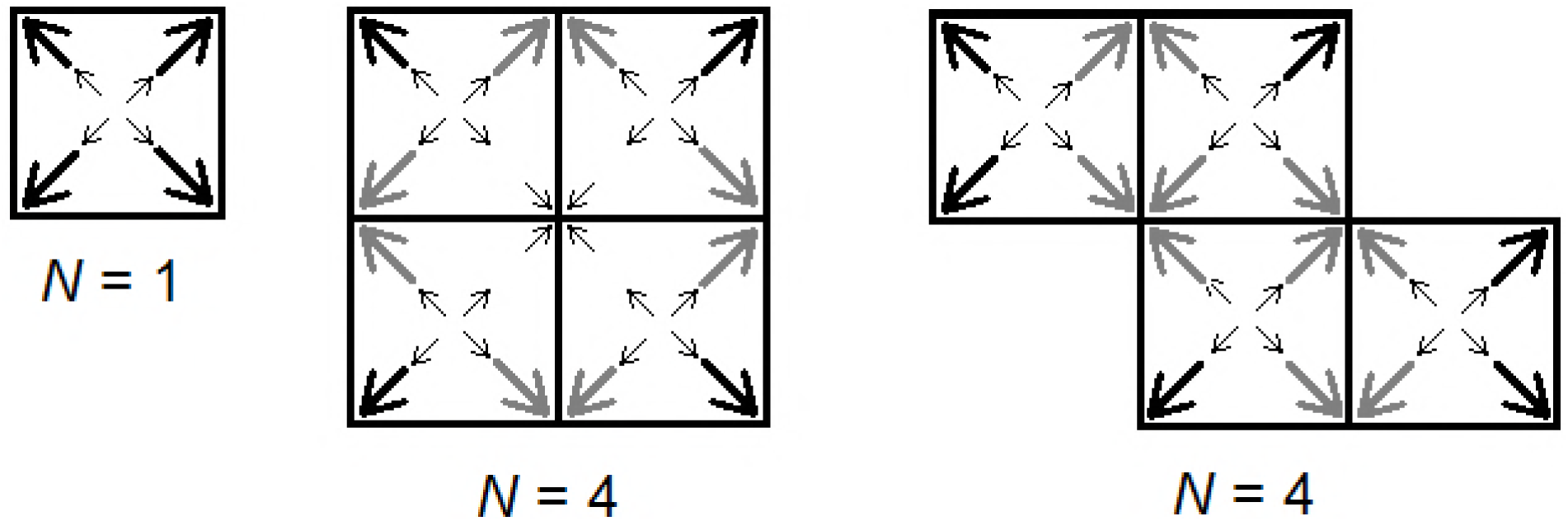
Force distribution in a single cell model (left), 4-cell square cluster model (center), and 4-cell non-convex cluster model (right). Thick black arrows indicate largest forces applied to the corners of the clusters; thick gray arrows represent intermediate forces applied at corners along the cluster edges; thin black arrows indicate small forces applied at the interior corners and near the cell centers.

Here 25 signifies the number of 5-min fluctuations over 2-h of Monte Carlo time.

For each cluster size *N*, we made 100 Monte Carlo simulations. For each simulation we computed the corresponding *CV_T_* according to Eq. 6. Finally, we plotted the mean *CV_T_* out of 100 simulations vs. *N*.

We first validated the results obtained from Eq. 4. We applied Monte Carlo forces whose means were 〈*F^e^*〉 = 8 units of force and 〈*F^i^*〉 = 1 units of force to the edges and interiors of the square and non-convex clusters, respectively, and obtained the corresponding *CV_T_* vs. *N* relationships. We found these relationships to be in an excellent agreement with the corresponding results obtained from Eq. 4 (Fig. Section S3 of Supplemental Information). Furthermore, when we applied Monte Carlo forces of the same mean value to both edges and interiors of the clusters, *CV_T_* attenuated with 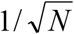, which was consistent with the prediction of Eq. 4 for the case where 〈*F^e^*〉 = 〈*F^i^*〉.

Since in clusters of living cells there are more than two types of traction forces acting on a cluster, we applied a broader range of forces in our Monte Carlo models. We assigned largest forces to the corners of the cell cluster in the range of 9-13 units of force; intermediate forces to the edges in the range of 3-9 units of force, and smaller forces to the interior cell-corners and to the center of each cell in the range of 1-3 units of force. The reason for this choice of force distribution is our previous observation of traction distribution in multicellular clusters of BAECs (unpublished data) and of fibroblasts (19) patterned on square-shaped islands. In these, the largest tractions were measured at the corners, and the smallest were measured in the interior of the cluster. In the non-convex clusters, at the corners where only three cells meet, we assigned intermediate forces (see Fig. 4, right panel). Initially, all forces were directed along the diagonals of the square cell units, away from the center of the cell. The forces were then equilibrated as using the least squares procedure, as in the earlier Monte Carlo simulations. Next we computed *CV_T_* for each of 100 simulations according to Eq. 6, and then obtained the mean *CV_T_* vs. *N* relationships for the square and the non-convex models (Fig. 8). We found that *CV_T_* decreased with increasing *N.* As found in the earlier model, the rate of decrease was somewhat greater in the non-convex clusters than in the square clusters, but smaller than the rate of the inverse square root dependence.

**Figure 8.**
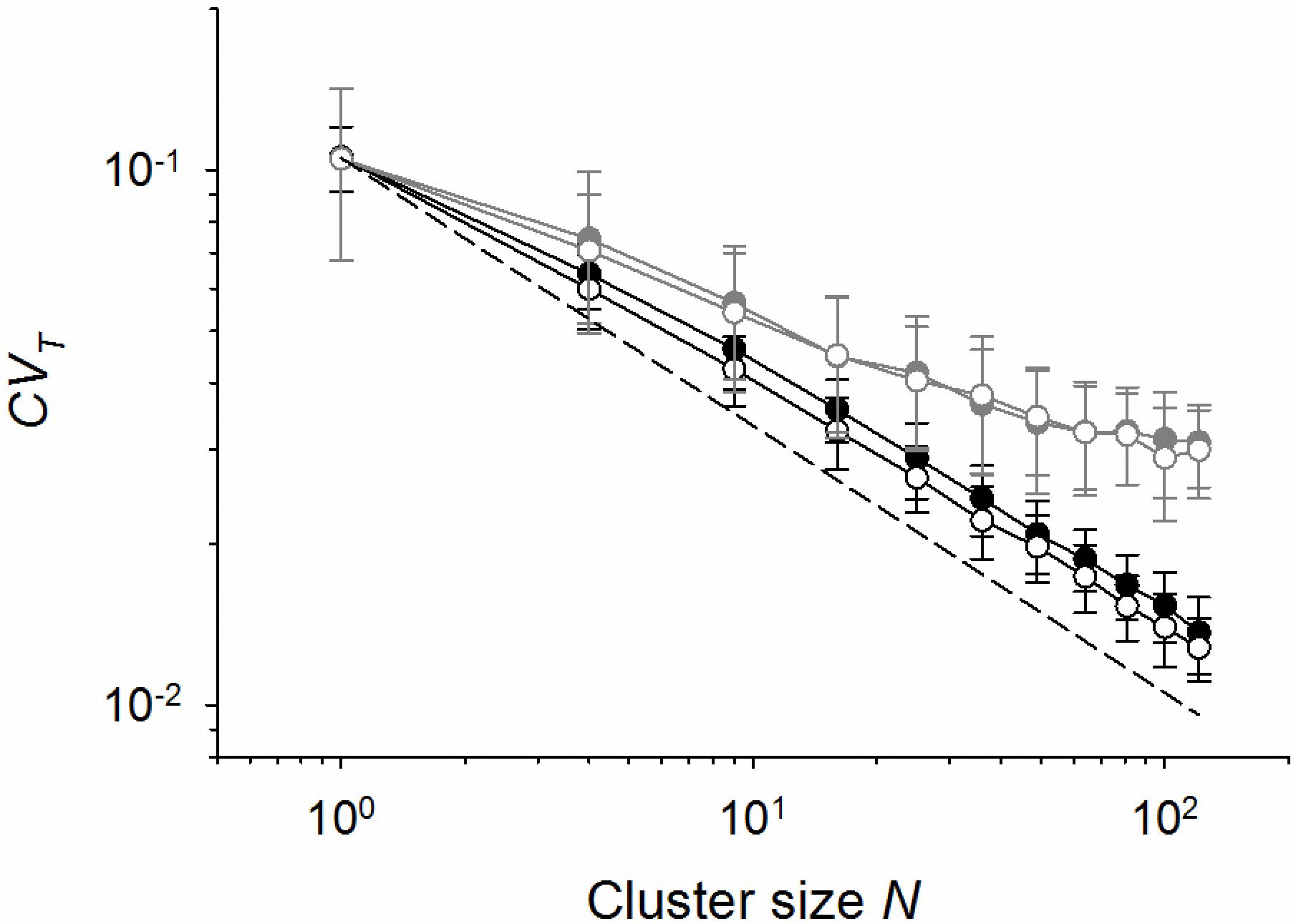
Coefficient of variation (*CV_T_*) vs. cluster size (*N*) relationship obtained with the cluster models with Monte Carlo forces (black) and with FA forces obtained from measurements in clusters of BAECs (gray). The solid circles correspond to the square clusters and the open circles correspond to the non-convex clusters. The dashed line corresponds to the inverse square root dependence, 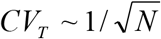. Data are means from 100 simulations ± SD.

#### Simulations with Measured Force Magnitudes

The force magnitudes assumed in the Monte Carlo simulation exhibit random fluctuations. On the other hand, measured FA traction forces in most cases do not exhibit random fluctuations (based on run test tables, two-tailed test, *p* < 0.05). To see to what extent this difference between Monte Carlo forces and measured FA forces affects traction field fluctuations of the cluster models, we applied measured forces of the same range of mean values as Monte Carlo forces to the models. From all FA forces that we measured previously in single BAECs and in clusters of BAECs (15), we randomly picked forces and assigned them to the corners (〈*F*〉 = 9-13 nN), edges (〈*F*〉 = 3-9 nN), and interior (〈*F*〉 = 1-3 nN) of the square and non-convex clusters based on the mean values of forces. Details of this procedure are given in the Supplemental Information (section S2). The forces were then equilibrated as in the case of Monte Carlo forces and the corresponding *CV_T_* was calculated according to Eq. 6. For each cluster size *N*, 100 simulations were made, the corresponding *CV_T_*s were averaged, and the mean *CV_T_* was plotted as a function of *N*.

We obtained that *CV_T_* decreased with increasing *N* at a lower rate than in the case of Monte Carlo forces (Fig. 8). There was virtually no difference between the *CV_T_* vs. *N* relationships of square clusters and non-convex clusters.

We repeated the above procedure using forces obtained previously from traction measurements on single cells and clusters of MEFs (16). In this case, the corner forces were in the range of 8-13 nN, the edge forces were 3-8 nN, and the interior forces were 1-3 nN. We found that attenuation of *CV_T_* with increasing *N* was smaller than in BAECs (Fig. 9). This is consistent with our previous observations that in MEFs traction field fluctuations were not significantly different between clusters and single cells, whereas in BAECs they were (16).

**Figure 9.**
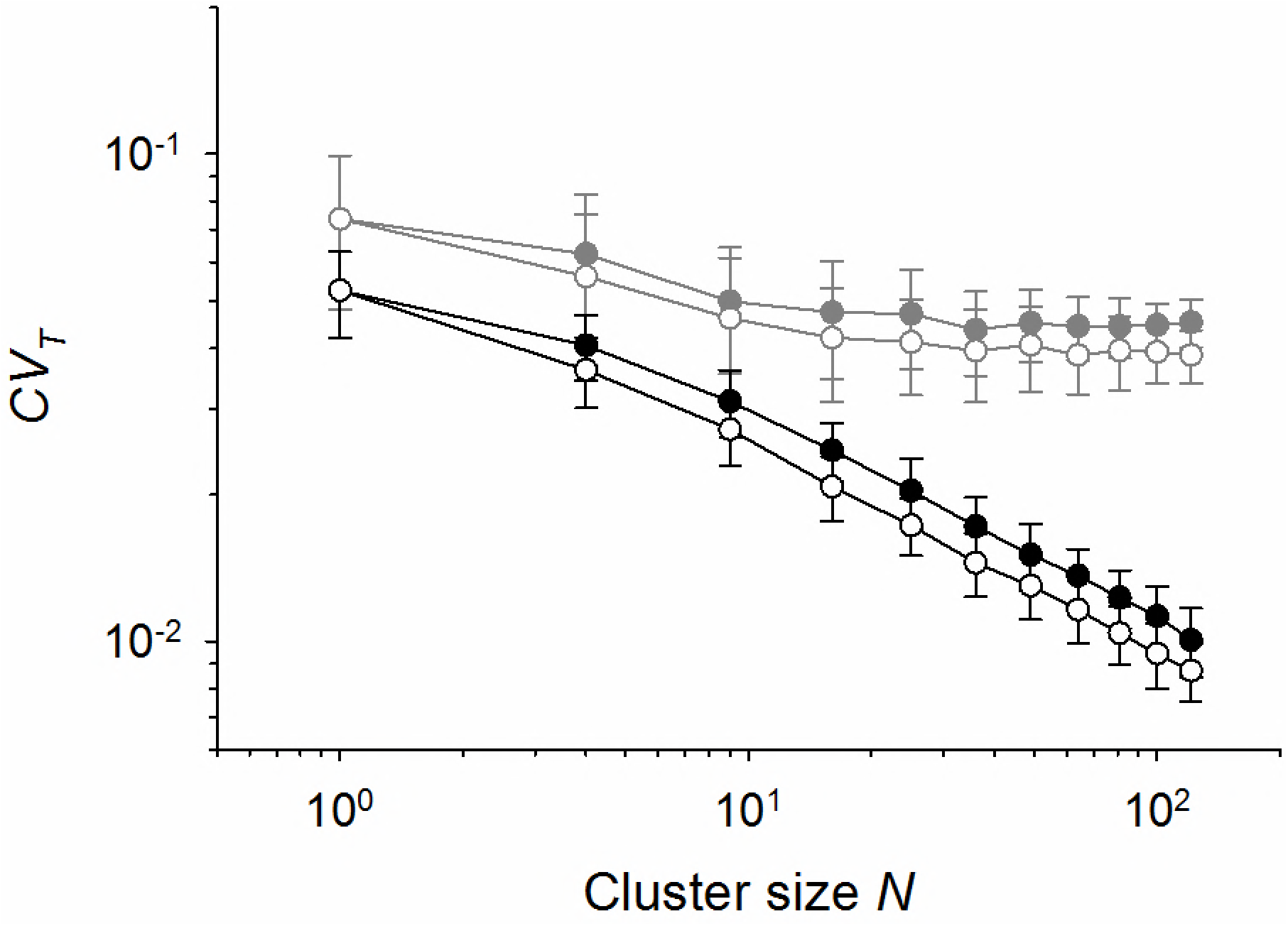
Coefficient of variation (*CV_T_*) vs. cluster size (*N*) relationship obtained with the cluster models with Monte Carlo forces (black) and with FA forces obtained from measurements in clusters of MEFs (gray). The solid circles correspond to the square clusters and the open circles correspond to the non-convex clusters. The dashed line corresponds to the inverse square root dependence, 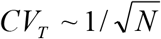. Data are means from 100 simulations ± SD.

## DISCUSSION

In this exercise, we analyzed how the distribution (i.e., edge vs. interior) and associated dynamics of FA traction forces, as well as how size and shape of cellular clusters affected variability of the overall traction field of multicellular clusters. These aspects of tensional homeostasis have not been previously explored and in that regard results of our analysis are novel. They provide a new insight into mechanisms by which tension is maintained stable in cells and tissues. We found that increasing cluster size and increasing edge to interior force ratio attenuated traction field fluctuations. Attenuation of fluctuations with increasing cluster size reflects the effect of statistical averaging. Attenuation of fluctuations with increasing edge to interior force ratio is a consequence of smaller variability of large edge forces vs. greater variability of small interior forces. However, increasing edge to interior force ratio tended to slow down the rate of attenuation due to statistical averaging. With regard to the cluster shape, the non-convex clusters exhibited a somewhat faster rate of attenuation of fluctuations of the traction field than the square, convex, clusters. This is a result of a greater fraction of edge forces relative to the interior forces in the non-convex clusters than in the square clusters. Together, the combined effects of statistical averaging, traction force variability, and of cluster shape, resulted in attenuation of traction field fluctuations with increasing cluster size at a rate that was slower than the rate of the inverse square root dependence.

Our analysis also showed that the traction field of Monte Carlo forces was attenuated with increasing cluster size at a faster rate than the traction field of experimental FA forces of similar magnitudes. This is an interesting finding considering that both Monte Carlo forces and FA forces had on average a similar *CV_F_* vs. 〈*F*〉 relationships. Thus, the observed differences in the traction field attenuation probably reflect the differences between the random fluctuations of Monte Carlo forces and non-random fluctuations of the FA forces. The non-random fluctuations of FA tractions reflect the dynamics of FA forces, which may rise, peak, or fall during the observation period. Such FA forces have higher values of *CV_F_* than Monte Carlo forces of the same 〈*F*〉, as can be seen from Fig. 1 vs. Fig. 7. Consequently, the overall traction field of the clusters with FA forces would exhibit greater values of *CV_T_* than the corresponding clusters with Monte Carlo forces.

We pointed out that model simulations obtained with FA forces measured in MEFs showed small changes in *CV_T_* with increasing *N*. This was consistent with our previous observations that traction field fluctuations of single MEFs were not significantly greater than that of multicellular clusters (16). In that study we used a different metric of the traction field, namely net traction moment, which is equal the sum of the dot product between traction forces and the position vectors of their respective points of application (see Section S3 of Supplemental Information). When we computed the net traction moment for our convex and non-convex models using MEF forces, we found that the corresponding coefficient of variation of the net traction moment remained virtually constant for different cluster sizes (Fig. S4, Supplemental Information).

Quantitatively, values of *CV_T_* obtained from the model with FA forces were underestimates in comparison with values that we obtained experimentally (15). A possible reason is that in our model we used only FA forces that remained finite (i.e., greater than 0.3 nN, which represents experimental threshold below which forces could not be distinguished from zero) throughout the 2-h observation period. However, a number of measured FA forces which were initially greater than 0.3 nN would decrease and stay below 0.3 nN, or decrease below 0.3 nN and then increase above 0.3 nN. Such forces had greater values of *CV_F_* than the forces that remained above the threshold value. Since these forces, as well as the forces that stayed above the 0.3 nN threshold, were included in calculations of experimental values of *CV_T_*, the obtained values tended to be greater than the values obtained from the model.

We needed to adjust the force field applied to the cluster at every 5-min interval in order to ensure that the forces satisfy static equilibrium. As a result of this adjustment, force vectors might change their directions and magnitudes. This, in turn, might have affected our estimates of *CV_T_*, especially in smaller clusters. Note that the same procedure was applied to the experimentally measured traction field (15,16), so if the force equilibration produced any bias in the estimated value of *CV_T_*, it was present in both modeling and in experimental results.

### Broder Impact

Endothelial cells and epithelial cells *in vivo* form continuous monolayers. Because of a very large number of cells and the absence of the edge vs. interior effects in those monolayers, tensional homeostasis in the monolayers is most likely primarily provided through statistical averaging of tensional fluctuations. However, injury or disease may disrupt monolayer integrity to create stress-free boundaries at the wound edges. Here, large traction forces may buildup, which, in turn, may cause loss of tensional homeostasis locally. For example, in the endothelium, maintenance of tensional homeostasis downregulates pro-inflammatory signaling (3). Stent implantation in coronary arteries, which destroys large endothelial segments, disrupts local stress transmission in the endothelium and creates conditions for breakdown in tensional homeostasis in the tissue surrounding the stent implant (9). Ensuing vascular inflammation can activate platelets and initiate thrombus formation, leading to life-threatening stent thrombosis. Regeneration of the damaged endothelial integrity is therefore essential for reestablishing tensional homeostasis, which provides a defense against thrombosis. Epithelial integrity is essential for tensional homeostasis in the epithelium and may provide a tumor suppressing mechanism. Disruption of epithelial integrity creates conditions for breakdown of tensional homeostasis in the epithelium, which is a hallmark of advanced breast cancer (2,6). The above examples illustrate the importance of understanding mechanisms which regulate and maintain tensional homeostasis. This knowledge may lead towards development of tools and procedures to improve wound healing.

### Conclusions

The analysis and models presented here gave us insights into mechanisms that might influence attenuation of traction field fluctuations in multicellular clusters. Besides statistical averaging, which we previously identified as a key factor that leads to attenuation of fluctuations of the traction field, our study revealed two novel factors that influence this attenuation, namely the difference in magnitudes of traction forces at the cluster edge and in the cluster interior, and dynamics of traction force fluctuations. The dynamics seems to be related to cell types and it can explain why in certain cell types, such as endothelial cells, cell clustering promotes tensional homeostasis, whereas in other cell types, such as fibroblasts, clustering has virtually no effect on homeostasis. Together, results of this study are important for understanding how cells maintain tensional homeostasis, which is critical for normal physiological function of biological tissue and for protection against various diseases.

## Author Contributions

J. Li carried out modeling and data analysis, and contributed to writing the manuscript and to figure preparation. P. E. Barbone contributed to the analysis of the coefficient of variation, to design of the Monte Carlo algorithm, and to writing the manuscript. M. L. Smith contributed to results interpretation and to writing the manuscript. D. Stamenović developed the concept, designed the research, drafted the manuscript and prepared figures.

## Acknowledgements

We thank Ms. Han Xue for providing experimental data. This study was supported by NSF grants CEBET 115467 (M. L. Smith) and CMMI-1362922 (D. Stamenović).

No conflicts of interest, financial or otherwise, are declared by the authors.

